# Leadership and information transfer in groups escaping a (simulated) threat in the wild

**DOI:** 10.1101/2023.07.05.547602

**Authors:** Akanksha Rathore, Bhavya Deepti Vadavalli, Vivek Jadhav, Kavita Isvaran, Vishwesha Guttal

## Abstract

Collective motion in many species is believed to be driven by predation, which is considered a crucial evolutionary force. However, there are limited studies on how information about predators spreads through a group, particularly in natural settings. Here, using high-resolution drone-based recordings, we analyze the collective escape dynamics of a group living species – blackbuck (*Antilope cervicara*), an Indian antelope – under the conditions of a simulated threat in their natural grassland habitat. Our analysis reveals that, in response to the simulated threat, group cohesion first increases, followed by a simultaneous increase in median speed and polarization. We also observe the emergence of a temporary leader-follower relationship during the collective escape. Interestingly, we found that the distance of individuals from the “predator” affected only their response time, not their influence on the group movement. The individuals furthest from the threat were the least likely to initiate movement and were uncoordinated with the group’s speed. Additionally, we discovered that the most influential individuals during the collective escape were the least likely to occupy front positions during group movement. Instead, our results indicate that individuals at the rear “push” individuals at the front during collective escape events. This contrasts with the typical notion of individuals at the front determining group movement and those at the back trying to catch up.

## 1 Introduction

Many group-living organisms produce fascinating patterns of collective motion. Mathematical and computational models show that collective patterns can emerge from simple local interactions among group members (Couzin, Krause, et al. 2003; Sumpter 2010; Vicsek and Zafeiris 2012; Deutsch, Theraulaz, and Vicsek 2012). For example, a tendency to be attracted towards nearby individuals can maintain cohesion whereas a tendency to align the direction of motion with nearby individuals leads to polarized group movement (Vicsek et al. 1995; Couzin et al. 2002; Jhawar et al. 2020). Group living and collective motion have important fitness consequences (Parrish and Edelstein-Keshet 1999; Murphy and Pitcher 1997; Bode et al. 2010; Hamilton 1971; Vine 1971; Watt, Nottingham, and Young 1997; Lack 1968; Grünbaum 1998). Predation, among various ecological factors, is recognized as a key driver for the selection of group living and collective motion (Krause and Ruxton 2002; Krause and Godin 1995). In fact, an experimental study involving actual predators and virtual prey has shown that predatory fish can adopt simple rules of local attraction and alignment among prey. (Ioannou, Guttal, and Couzin 2012). Anecdotally, studies have described flash expansion and fountain effect patterns when prey groups swirl away from predatory attacks (Murphy and Pitcher 1997). While exploring this phenomenon, it is crucial to reiterate that the information about a predatory threat is likely available locally to a few individuals in the group and is then rapidly transmitted to the entire group. Only a few empirical studies have looked at the fascinating phenomenon of collective escape dynamics, linking local rules of organisms to group-level patterns, either in a laboratory or natural conditions (Treherne and Foster 1981; Ballerini et al. 2008a; Sumpter 2006; Handegard et al. 2012; Sankey et al. 2021; King et al. 2012; Poel et al. 2022; Papadopoulou et al. 2022b; Papadopoulou et al. 2022a; Ioannou et al. 2019; Marras and Domenici 2013; Jolles et al. 2021; Ioannou et al. 2023; Lambert, Herbert-Read, and Ioannou 2021). To fill in this major gap, this paper focuses on investigating collective escape dynamics and the intricate underlying mechanisms that give rise to such dynamics in blackbuck herds in their natural habitat.

We briefly summarise some of the empirical studies on collective escape dynamics. While some studies have focused on group-level collective escape patterns, others have looked at the effects of predatory threats on local interaction rules. Handegard et al. 2012 did an impressive study on how group hunting by a predatory fish (spotted sea trout, *Cynoscion nebulosus*) affects collective escape behaviour in a schooling prey (juvenile Gulf menhaden, *Brevoortia patronus*) in the natural estuarine and near-shore ecosystems. They used SONAR-based high-frequency imaging to track the movement of individuals and groups. Focusing on group-level patterns, they find that information transfer among group members of prey fish increased with group size. However, coordinated hunting by predators led to an increased frequency of splitting among prey schools, thereby reducing the efficiency of information transfer among prey. In Romenskyy et al. 2020, again focusing on group-level patterns, using a controlled experiment that involved both the prey (*Pseudomugil signifer*) and live predators (*Philypnodon grandiceps*), authors found that prey shoals were present on the surface of the water in a carpet-like structure whereas they formed a sphere-like structure under-water (Romenskyy et al. 2020), possibly minimizing the overall surface area available for a predatory attack.

A few experimental studies have looked at how schooling prey modify their local interaction rules in simulated predatory threats (Herbert-Read et al. 2015; Herbert-Read et al. 2017). By simulating a predatory stimulus on a schooling fish (Pacific blue-eyes, *Pseudomugil signifer*), Herbert-Read et al. 2015 showed that changes in the velocity of a small number of individuals could spread across the entire group, in the form of high-density escape waves. Furthermore, a study on shoals of Trinidadian guppy (*Poecilia reticulata*) showed that fish modify the local behavioural rules of interactions (Herbert-Read et al. 2017; Storms et al. 2019). For example, fish attract each other at shorter distances than in the predator-free condition. However, interestingly, their alignment rules remain unchanged.

In fission-fusion societies – like our study system blackbuck – groups merge and split relatively frequently. Therefore, stable hierarchical relationships are unlikely to exist and sustain. Thus, the process of collective decision-making is likely to be egalitarian rather than despotic. However, individuals may still differ in their responses due to inherent heterogeneity in individual behaviours. Such variations may result in a context-dependent hierarchy or temporary leaders who “lead according to need” (Conradt and List 2009). Therefore, the key to unravelling collective dynamics lies in identifying individuals’ behavioural rules and how information flows between individuals. However, we currently know little about both, the way animals interact with each other during predatory threats and the ensuing group-level patterns of collective escape, especially in the natural conditions.

Studying collective motion and in particular predatory collective escape in the wild, however, is challenging for several reasons. First, predatory events are rare and difficult to observe. Secondly, collecting data on such events where we can track multiple individuals, preferably all individuals, of the herds can be an arduous task. Third, even if such data are available, analysis and interpretation of the response of a large number of individuals of herds themselves pose many challenges.

In this study, we aim to understand how individual behaviour affects the collective response of animal groups to predation-like events in blackbuck herds. We overcome many of the challenges associated with such a study via a combination of (i) design of a novel and relatively noninvasive simulated threat, (ii) a noninvasive UAV-based high-resolution video recording of the collective motion and escape response of the herds, (iii) implementation of the state-of-the-art animal tracking techniques from such high-resolution videos and finally, (iv) analyses of the tracked trajectories to investigate the patterns of collective escape using techniques from the physics and social network science literature.

We expect that in blackbuck herds, which exhibit ephemeral groups due to fission-fusion dynamics, decision-making is egalitarian. More specifically, we address our questions in the following order: We first look at how group patterns change during the escape response. We then investigate how these group patterns relate to individual-level behaviours. There are two reasons to focus first on group-level patterns: First, there are standard and relatively easy-to-use metrics to measure group dynamical properties. Secondly, but importantly, measurement of group-level patterns can provide insights into individual-level behaviours (Rathore, Isvaran, and Guttal 2023; Ballerini et al. 2008b; Jhawar et al. 2020; Jhawar and Guttal 2020). We then explore the role of individuals in shaping group response by identifying response initiators and individuals who exert disproportionate influence over group movement decisions.

## 2 Data Collection

We studied the escape behaviour of blackbuck (*Antilope cervicapra*) herds in response to induced perturbations (simulated threat). Detailed information about the study species and the site can be found in SM section 1. To obtain a comprehensive understanding of the collective behaviour of these herds, it was necessary to observe all members and their interactions simultaneously. Recent advances in aerial imagery techniques have made it possible to obtain high-resolution movement data without the need to capture and tag animals. With the necessary permissions and ethical approvals from the Gujarat state forest department and our institutional animal ethics committee, we used DJI Phantom-4 unmanned flight vehicles (as shown in Figure 1) to record herd behaviour in aerial videos. Data collection took place between February and March 2017, during which we recorded four videos each day for different herds: two in the morning and two in the evening, with a duration of 10-15 minutes each (limited by battery life). As predator attacks and the group’s escape response are brief events, a 10-minute duration is sufficient to observe fine-scale movement behaviour.

**Figure 1:**
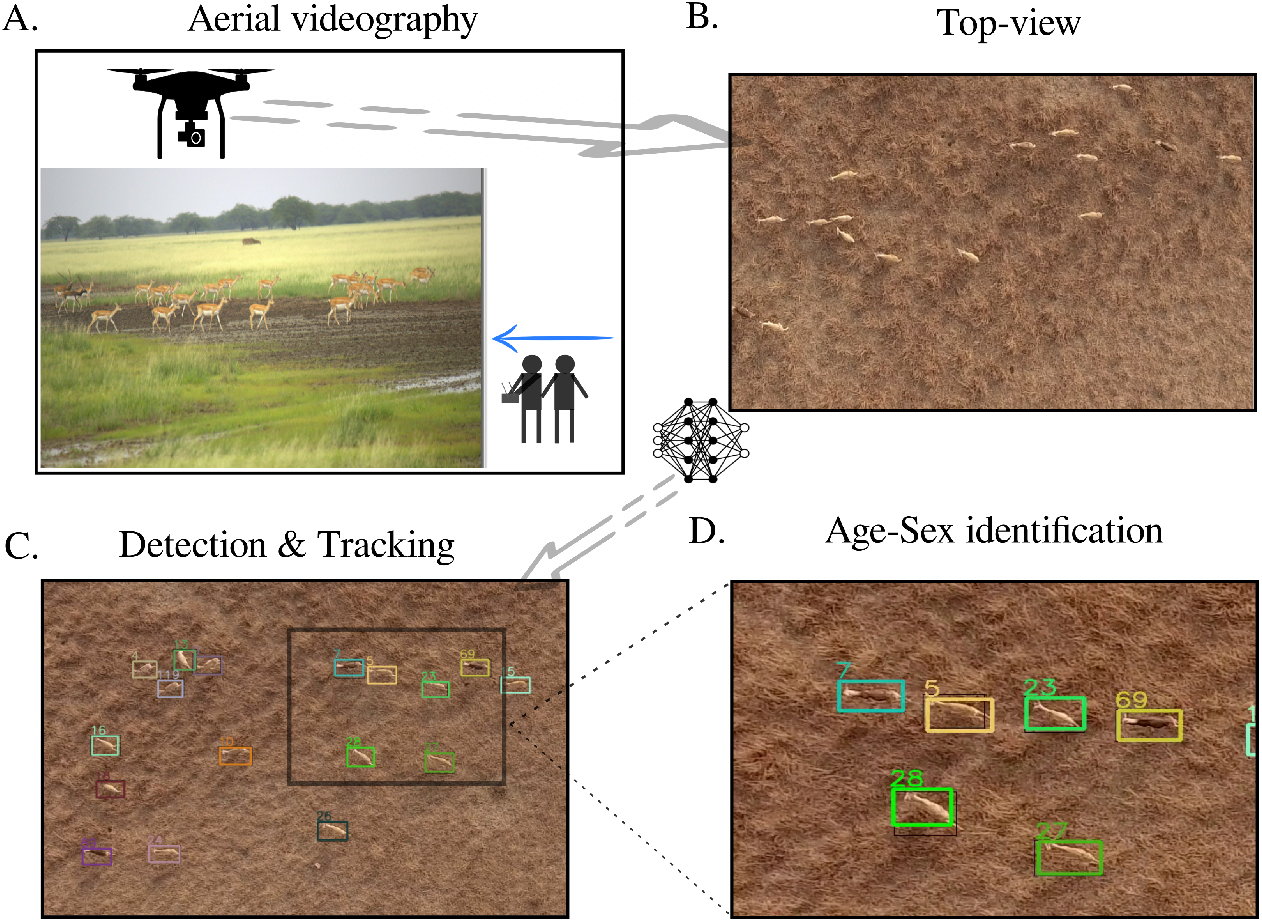
A. We record group behaviour by flying a drone over the group, to introduce a threat-like perturbation one of the observers walks towards the herd, B. We get top-view videos in which all the individuals and their movements can be observed simultaneously, C. To get the spatial positions and trajectories of the individuals in these videos, we perform machine-learning-based detection and training, D. The high-resolution videos allow us to zoom in and identify the age-sex category of the individuals.

To overcome the challenge of observing predation events in the wild, we created a mild threat-like scenario or perturbation. It is well-known from the previous studies on blackbuck in these areas that these animals typically avoid human proximity (Ranjitsinh 1982; Jhala and Isvaran 2016). We utilized this knowledge to design a relatively mild and noninvasive form of perturbation. During the recording, one of the experimenters would walk towards the herd at a normal pace approximately halfway through (5-7 minutes). As expected, the blackbuck herd responded by moving away from the human experimenter (refer to Figure 1-A). Once a response was induced, we ceased our approach and continued recording their collective movement response. It is worth noting that blackbuck in Velavadar National Park are used to human presence, as the park is a well-known destination for blackbuck and grassland enthusiasts from around the country. The forest department takes various measures to maintain the watering holes in the park. Moreover, poaching and hunting are rare in the surrounding areas. Hence, it is likely that blackbuck consider human presence or movement as a mild perturbation or nuisance.

We conducted approximately 75 events in various types of habitats (grassland, shrubland), and processed the resulting videos using the method outlined in Rathore et al. 2023. This process extracted the movement trajectories of all individuals present in the videos. Although the process was automated, the complexity of the background resulted in inaccuracies in track linking. To correct these errors, we manually connected the lost IDs. For our analysis, we selected 26 videos that were recorded exclusively in the grassland habitat.

## 3 Methods and Results

### 3.1 Four phases of group behaviour in collective escape dynamics

Upon visual inspection of the videos, we identified four qualitatively different types of behavioural states during the experiment. These states are also distinct in terms of the group’s speed, the time taken to initiate the response, and the group’s structure during the response phase (as depicted in Figure 2-A).

**Figure 2:**
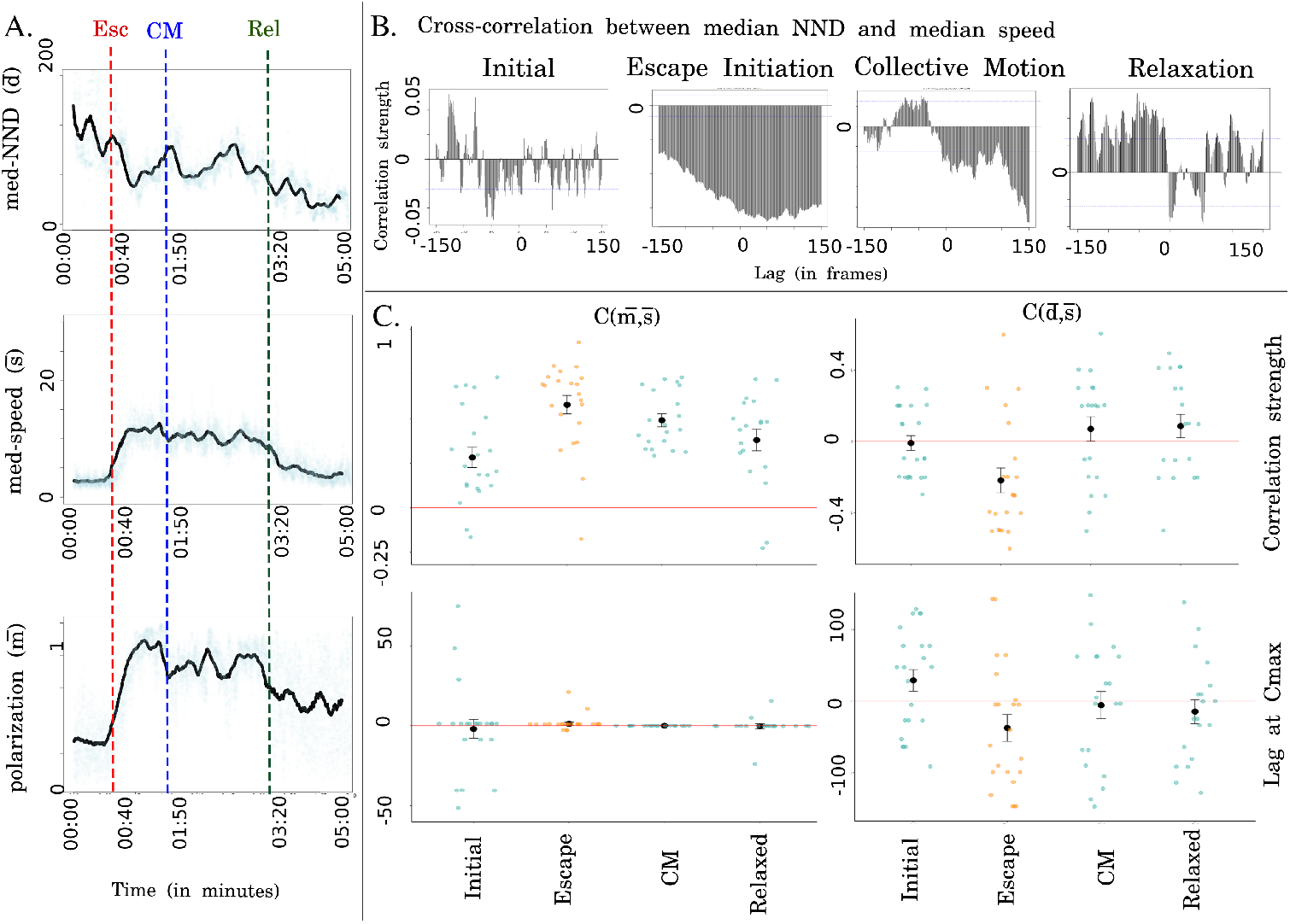
A. Time-series of median nearest neighbour distance, median speed and polarization metrics calculated for every frame of a 5-minute video, time-stamps for the visually identified movement phases are marked with coloured dotted lines B. Cross-correlation function (between median NND and median speed) calculated for the escape phase for one of the videos C. Pooled results across 26 videos for the cross-correlation between polarization & median speed and median NND & median speed.

1. **Initial unperturbed** - This is the initial phase where the group behaviour is recorded without any external disturbance or perturbation. This phase serves as a baseline for the subsequent phases.
2. **Escape initiation** - This phase corresponds to the herd’s response initiation behaviour, where individual members of the group become aware of the approaching threat and begin to move away.
3. **Collective response** - In this state, all individuals within the group have initiated the escape response and move together in a seemingly coordinated manner.
4. **Relaxation** - This phase occurs after the escape response when all the individuals stop moving and resumed their normal activities, similar to the initial phase before the perturbation event.

To better understand the collective escape dynamics observed in these videos, we quantified the group-level patterns displayed by the animals. These patterns are commonly used in the collective behaviour literature to infer the degree of local attraction and alignment tendencies within animal groups (Sumpter 2006; Herbert-Read et al. 2017; Herbert-Read et al. 2011). To compute these quantities, we first need to detect and track the animals from high-resolution drone videos. We do so by employing the package MOTHe (Rathore et al. 2023), which employs a convolution neural network-based machine learning approach for animal detection in heterogeneous backgrounds. The visual animal tracking is based on the Kalman and Hungarian algorithms. Based on the tracks obtained, we measured the following for each frame of the video:

1. **Median nearest neighbour distance (median NND or** 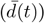: We first compute the distance to the nearest neighbour (denoted by *d*_*i*_(*t*) at time *t*) for a focal individual *i*. We then compute the median of the nearest neighbour distances (median NND or 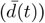. This metric can be thought to represent group cohesion with a larger value implying a lower group cohesion and vice-versa. We also note that group cohesion typically arises from local attraction tendencies among individuals in a group (but cohesion may also be a consequence of local direction alignment). Thus, group cohesion may be used to infer the strength of local cohesive tendencies among group members.
2. **Group polarization** 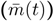 is the vectorial sum of individuals’ orientations (*v*_*i*_(*t*)) divided by the number of individuals in the group (Vicsek et al. 1995; Vicsek and Zafeiris 2012),

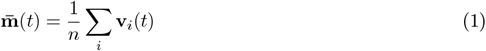

Group polarization quantifies the degree of direction alignment among group members. It is often referred to as the order of the collective motion. In models of collective motion, increasing the tendency of animals to align locally with their neighbour’s direction enhances group polarization (Couzin et al. 2002; Ioannou, Guttal, and Couzin 2012). Therefore, conversely, group polarization can be used to infer the strength of local alignment tendencies among animals. For this work, we focus on the scalar quantity 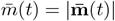.
3. **Median speed of individuals** (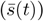: We compute the speed of all individuals, denoted by *s*_*i*_, between consecutive frames based on the tracking of animals and then compute the median value for each frame. Computational models of collective motion typically assume a constant speed (Vicsek et al. 1995; Couzin et al. 2002). However, here we expect that this quantity could be a crucial metric to capture the immediate response of herds to the perturbation (and hence predatory threat).

In Figure 2-A, we show the time series (after smoothing i.e. averaging for 300 data points that correspond to 10 seconds in our videos) for the above three metrics for one video for a duration of five minutes around the time of perturbation (See SM Figure 1 for actual time-series).

### 3.2 Relationships between different phases of group responses via cross-correlation function

Next, we examine how these three metrics interact to investigate the dynamics of the group’s response and to deduce the local response rules of blackbuck. To accomplish this, we employ the computation of cross-correlation functions (CCF) between the corresponding time series, which is elaborated upon below.

#### 3.2.1 Cross-correlation functions

We assess the temporal cross-correlations among the following three time-series: median nearest-neighbour distance 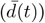, group polarization 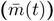, and median speed 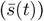. Cross-correlation serves as a metric for gauging the similarity between two time-series as a function of the temporal displacement of one relative to the other.

Formally, cross-correlation function between two time series *x*_*i*_(*t*) and *y*_*i*_(*t*) is defined as

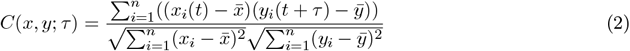

where *n* is the number of time points over which data is available and 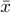 and 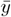 are sample means of *x*_*i*_ and *y*_*i*_ time series.

From the cross-correlation function (CCF), we identify the *time lag* at which pairs of time-series are *maximally correlated* (Nagy et al. 2010). We use the value of this time lag (denoted *τ*_*m*_) and the strength of correlation at this lag to infer the group response rules. We also note that a significant correlation has a value greater than 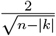, n is the number of observations and k is the time lag.

The interpretation of the relationships is as follows:

##### Cross-correlation value

A positive cross-correlation value indicates that an increase in one value of a time series corresponds to an increase in the other value of another time series (Nagy et al. 2010).

##### Sign of lag

The sign of the lag indicates the direction of the relationship, indicating which series changes first. Specifically, a positive lag between *x* and *y* suggests that a change in *y* occurs before a change in *x*.

#### 3.2.2 Relative role of the attraction and alignment tendencies

The metrics that we measure at the group level represent the local tendencies of attraction and alignment within a group (Giardina 2008; Lukeman, Li, and Edelstein-Keshet 2010). Therefore, to gain insight into the relative significance of these tendencies, we analyze the lead-lag relationship in the time-lagged correlations. In Figure 2-B, we present the output of the cross-correlation function for the nearest-neighbour distance (NND) and speed, using an example video (refer to SM Figure 2 for all three pairs of group properties). We calculate the values of *τ*_*m*_ and the corresponding cross-correlations at *τ*_*m*_ for all 26 videos. The results of these calculations are depicted in Figure 2-C.

Through the analysis of these videos, we observe a positive correlation between group polarization 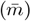 and speed 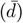 during all movement phases (Figure 2-C A, upper). However, we do not identify any causal or directional relationship as the time lag *τ*_*m*_ is 0 (Figure 2-C, lower). This suggests that group polarization and speed increase simultaneously during all phases of group behaviour, including the escape and coordination bouts within the collective escape. Following that, we observe that the median cross-correlation between nearest-neighbour distance (NND) and speed 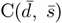 is negative during the escape and coordinated bout phases (Figure 2-C, right column). Additionally, the maximum correlation occurs at a negative time lag for the escape initiation phase. Based on these findings, we deduce that increased polarization and higher cohesion are correlated and occur simultaneously during the escape phase. However, an increase in group cohesion precedes group polarization during the escape initiation. From these observations, we infer that in response to a threat or disturbance, the group first becomes cohesive, followed by a simultaneous increase in speed and alignment. Consistent with the aforementioned analysis, we find that the median cross-correlation between NND and polarization 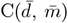 is negative for the escape and coordinated bouts (Figure 2-C). The associated lag is negative only for the escape initiation phase (refer to SM Figure 3). The pattern of the cross-correlation function (CCF) 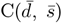 is expected to be similar to that of 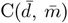 because median speed and group polarization are strongly positively correlated.

**Figure 3:**
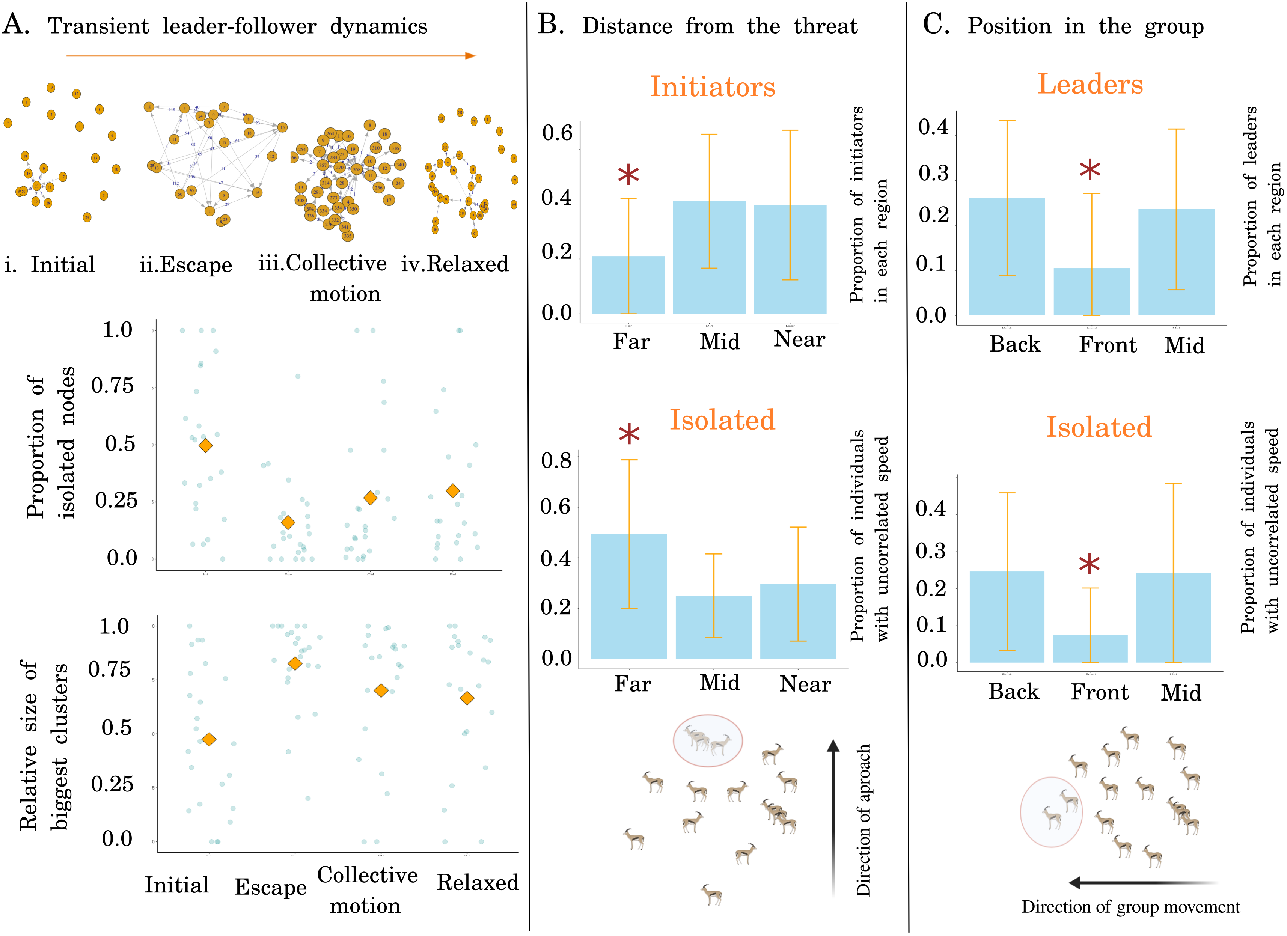
A. The social network structures were constructed by computing cross-correlations in speed between all pairs within a group. An edge represents a cross-correlation strength surpassing 0.6, and the edge weight indicates the time lag. This figure displays the percentage of individuals without any connections and the relative size of the largest clusters across all videos. B. The distribution of individuals initiating the escape response is based on their distance from the threat. The second figure presents the distribution of individuals whose speed is uncorrelated with group movement, categorized by the distance from the threat. C. The distribution of individuals exerting the greatest influence on group movement, corresponding to their spatial position in the direction of group motion. The second figure exhibits the distribution of individuals whose speed is not correlated with group movement at different spatial positions within the group.

Put together, we infer that while responding to a threat/perturbation, the herd first becomes cohesive; and then speeds up and orients simultaneously. In terms of individual-level interactions, we suggest that this could be due to a greater tendency to stay with others when threatened/frightened (King et al. 2012). Therefore, increased local attraction tendencies among group members could be the primary driver of the patterns at the group level.

### 3.3 Individual-level response

We now examine the local response and the role of individuals in shaping the collective escape dynamics of blackbuck herds. Specifically, we investigate the role of heterogeneity in groups – arising from age, sex and spatial position – in the collective escape dynamics of blackbuck herds. First, we examine how the information of threat propagates within the herd. To do this, we identify which individual of the herd initiates the response and how does this response spread to the entire herd. We then identify whether some individuals are more influential than others in determining the group response. See Glossary for the definitions of initiators, leaders and information transfer in the context of our study.

## Box: Glossary

**Information transfer** – Change in the speed of one individual followed by the change in speed of another individual (movement)

**Response initiators** - Initial individuals (one to five) to respond (in terms of change in speed)

**Laggers** - Last to respond (in terms of change in speed)

**Influential individuals (or leaders)** – Those whose movement appears to be followed by others (in terms of leading in cross-correlation patterns)

**Isolated** - Individuals performing temporally uncorrelated movement

### 3.3.1 Response initiators

We identify the response initiators, i.e. individuals responding first to the approach, by quantifying changes in the speed of each individual in the herd. We first calculate the speed of all the individuals for every frame of the video by calculating the change in the position from *n*^*th*^ to *n* + 10^*th*^ frame. Then, we identify the points in time where the speed for each individual exhibits an appreciable change (referred to as ‘jumps’) after smoothing the time-series (SM Figure 5). We rank individuals according to the time we see the first jump in the speed and then select the top five responses; we term them as “initiators “. SM Figure 5 shows the initiators marked on one video frame during the response. We repeat this process for all the videos and note down the initiators’ IDs (top five responses) for each video.

After identifying the response initiators, we examine if the response initiators are governed by either intrinsic or extrinsic factors, i.e. age/sex and spatial positions. For this, we assign age-sex class categories, distance from the threat and spatial position during the collective motion as described in section 2.3 to all the identified response initiators. We then investigate whether individuals occupying a particular spatial position (near, mid, far) are more likely to respond early or late. For this, we calculate the proportion of individuals responding from each region. Figure 3-B depicts the proportion of initiators from each region. We observe that response initiators were likely to be in the near and mid-region and least likely to be from the further region. We also observe that the individuals that are furthest away from the threat don’t correlate their speed/movement with other group members (see “Isolated” in Figure 3-B).

Next, we examined whether individuals of a particular age/sex group are more likely to be initiators or laggers (SM Figure 4-A). We calculated the proportion of response initiators and laggers from each age/sex category. We observe that females are more likely to be the response initiators however the effect size is too low.

### 3.3.2 Leader-follower dynamics

After identifying the response initiators, we examine whether leader-follower relationships exist (or emerge) during the escape response. Given the high rate of fission-fusion dynamics in blackbuck herds, stable hierarchical structures are unlikely, and we expect the dynamics to be self-organised and decision-making egalitarian. However, as observed in many other species (Nagy et al. 2013), the hierarchical structure can be context-dependent and even for self-organising groups there might be some individuals who exert disproportionate influence over group decisions (Conradt and Roper 2005; Conradt et al. 2009). Therefore, we examine the dynamics of groups in all four movement phases and explore the possibility of leader-follower dynamics.

We assume that the speed of movement of blackbuck contains information and thus we quantify information transfer in terms of change in speed (see Box). Therefore, we calculate pairwise cross-correlations in the time series of individual speeds to identify lead-lag relationships. We build a network of lead-lag relationships for the entire group for each video. To identify only the significant relationships, we arbitrarily choose a threshold on the correlation strength (>0.5) to build this network. Since we expect that any such relationships may rapidly change, we construct such lead-lag networks for the four-movement phases separately in each video.

See SM Figure 6-A for a cartoon of two hypothetical networks for a group of three members. A network of the type SM Figure 6-A-i indicates that information flows from A to C to B whereas a network of the type Figure 6-A-ii suggests that information flows from A to C to B, as well as directly from A to C. In both cases, A will be the most influential individual (or a *leader*) whereas C and B form *followers*. Figure 3-A shows pair-wise speed correlation networks for one video. We find that there are no lead-lag relationships during the initial and post-response phases (A.i and A.iv). However, connections emerge during the “Escape” and “Collective motion” movement phases (A.ii and A.iii) indicating the emergence of leader-follower dynamics during the predation-escape response.

We repeat this process for all the phases of 26 videos. If lead-lag relationships emerge during collective escape response, we should see an increase in the number of leader-follower pairs in the network during the escape and coordinated motion phases in all the videos. More specifically, we calculate the proportion of connected individuals and solitary individuals. We expect that the proportion of connections must increase while that of solitary individuals must decrease when leader-follower relationships exist. We also calculate the size of the biggest cluster of connected individuals and show it as a fraction of the total individuals in the group. As observed in the second graph of Figure 3-A, the proportion of connected individuals increases during escape response phases.

Indeed, we observe that during both the escape and collective motion bouts, the median proportion of unconnected nodes decreases and cluster size increases, indicating the emergence of leader-follower dynamics during the response (Figure 3-A).

### 3.3.3 Influential individuals

Next, we ask if some individuals exert more influence over the group movement than the rest, referred to as *Influential individuals or Leaders*, during the escape response. As shown in Figure 3, we use directed graphs to exhibit the relationship between the individuals, it’s because we represent the flow of information through lead-lag relationships in these graphs. Further, we calculate the strength of influence using the number of inward edges and edge weight that signifies the lag. This information helps us in identifying the nodes which spread information rapidly and hence influence the network. In our context, it would be equivalent to the individuals who spread the information fastest to the maximum number of group members. In SM Figure 5-C, we show a network colour coded for the degree of influence. This process was repeated for all 26 videos and in each video, the top five individuals were identified as influential individuals (or leaders).

We then examined whether the response initiators in these groups are the same as the influential individuals. Based on the analysis of 26 videos, we find that response initiators were not necessarily the most influential individuals. Next, we investigate whether individuals occupying a particular spatial position (near, mid, far) are more likely to exert a disproportionate influence over others during the escape response. For this, we calculate the proportion of individuals responding from each region. SM Figure 7-B depicts the proportion of influential individuals from each region in terms of the distance from the threat. We don’t find any discernable pattern here. Next, we examine whether individuals who exert more influence occupy a particular spatial position in the group with respect to the movement direction of the group. We expect leaders to occupy the frontal positions in the group. So, we calculate the proportion of leaders from all regions (front, mid, and back). Figure 3-C depicts the proportion of influential individuals from each region. We observe that contrary to our prediction, leaders or influential individuals are least likely to be at the front of the group. It is also interesting to note that the individuals at the front of the group are least likely to perform uncorrelated movement, this signifies that the individuals at the front are followers during the group response.

## 4 Discussion

Only a few studies have previously investigated the collective escape dynamics at high spatial and temporal resolution in their natural conditions. In the estuarine predator-prey system studied in (Handegard et al. 2012), SONAR was used to record trajectories of predator and prey at high-resolution. However, researchers had no control over the dynamics of predatory attacks on schooling prey. Tracking each individual for the entire duration was not feasible due to the presence of multiple groups over the video frame. Therefore, one could only study the group-level patterns of collective escape response. In our case, we used a simulated threat on blackbuck herds in open terrestrial habitats, allowing us to observe how the group activity changed from an initial foraging state. We could track how the response was initiated and spread in the entire group, while also tracking each individual of the herd. This allowed us to decipher not only group-level patterns but also make inferences about individual-level roles in the group response.

In summary, we argue that individuals exhibit a typical set of collective decision rules in case of threat-like perturbations in the fission-fusion blackbuck society. Despite variation in the group composition (age, sex structure and behaviour) of blackbuck herds, there are various characteristic features of collective escape dynamics across all groups we studied. However, a vital factor which may influence the group dynamics is habitat structure. So far, we have analysed only the videos which were recorded in open grassland habitats, and we may expect that the decision rules and group dynamics to be different for a patchy habitat i.e. shrubby/woody areas. It would be exciting to look into the dynamics of these groups in other types of habitats that could affect visibility and information flow. It is also interesting to investigate the generality of our results in other fission-fusion societies.

The collective escape dynamics of group-living animals in response to predation is an important area of study that has implications for both ecology and evolution. Our study used high-resolution drone-based recordings to investigate the collective escape dynamics of blackbuck in response to a simulated threat. Our analysis revealed several important insights into the collective behavior of blackbuck and the ecological implications of these results are discussed below.

We found that blackbuck prioritize group cohesion over individual escape during collective escape events. This suggests that group cohesion is an important mechanism for survival in the face of predation. Maintaining group cohesion can reduce the likelihood of individual capture and increase the chances of group survival. Our findings also highlight the importance of considering social behaviour in studies of predation risk.

Next, we observed the emergence of a temporary leader-follower relationship during the collective escape of blackbuck. This finding has significant implications for understanding the evolution of leadership in social animals. Temporary leadership structures that emerge during collective escape events may facilitate group survival by increasing coordination and reducing individual-level risk. Future studies could investigate the factors that determine individual differences in leadership ability and the mechanisms underlying the emergence of temporary leadership structures during collective escape events.

Furthermore, we found that the distance of individuals from the predator affected only their response time, not their influence on group movement. This finding challenges the commonly held notion that individuals at the front of the group determine group movement, while those at the back try to catch up. Instead, our results suggest that individuals at the rear “push” individuals at the front during collective escape events. These findings have important implications for understanding the role of individual differences in social behaviour in shaping group dynamics. Future studies could investigate the ecological implications of social behaviour in other species and the mechanisms underlying the emergence of temporary leadership structures during collective escape events.

## 5 Conclusion

In this study, we characterised the collective escape dynamics of blackbuck herds to a simulated perturbation/threat event. Our analysis of group dynamics revealed signatures of four different phases in the group activities: a baseline phase, an escape initiation phase, a collective motion phase and a post-response phase. We demonstrate that the blackbuck herds respond to threats by first increasing group cohesion, and then increasing group polarization alignment and group speed (both of these actions happen simultaneously). Furthermore, we find that when leader-follower relationships exist, they are short-lived and not persistent. We infer that collective escape dynamics in fission-fusion grouping animals are likely driven by egalitarian decision-making processes. We also identify the response initiators and the most influential individuals in this collective escape response. We expected these individuals, called response initiators, to guide group movement during the response. However, we observe that the most influential individuals are rarely the same as the response initiators. This result suggests that anyone can emerge as a leader in these groups. The probable reason for this would be that an individual’s influence is guided by other factors (intrinsic motivation or habitat cues) rather than access to information about predatory threats.

Further, we delved into the role of individual heterogeneity (owing to either age, sex or spatial positions of individuals) in shaping the response of individuals and consequently collective escape dynamics of blackbuck herds. Our results indicate, as intuitively expected that the individuals who are closer to the approach of a potential threat are more likely to be response initiators, and individuals further away from the approach are more likely to respond last. We also find that females are more likely to be the response initiators, whereas breeding males are likely to respond last. Finally, we find that breeding males and females are more likely to be influential individuals than juveniles or other males. However, these relationships are not statistically significant therefore it needs to be investigated further. We also find short-term leader-follower dynamics during which some individuals exhibit disproportionate influence over the group. Further analysis revealed that the individuals at the back of the group are likely to be influential individuals or leaders.

Overall, our study provides important insights into the collective behaviour of blackbuck in response to a predator threat. These findings highlight the importance of considering social behaviour in studies of predation risk and the evolution of leadership in social animals. Our study also provides a framework for studying group-living animals in the wild using non-invasive technological tools and novel data analysis methods to infer social interactions.

## 6 Author contributions

AR, KI and VG conceptualised the study. AR, VJ, KI and VG contributed methods. AR collected data and conducted analyses. BV assisted in data extraction and analyses. VJ assisted in modelling the escape behaviour. AR and VG wrote the manuscript with inputs from others.

## 7 Acknowledgements

We thank our TEE-LAB members at CES for many fruitful discussions. VG acknowledges the DBT-IISc partnership program and DST-FIST. AR acknowledges MHRD for the PhD scholarship and CRG-CASCB joint grant by the Department for the Ecology of Animal Societies at the University of Konstanz/Max Planck Institute of Animal Behavior and the Centre for the Advanced Study of Collective Behaviour, University of Konstanz.

