## Supplementary material for "Leadership and information transfer in groups escaping a (simulated) threat in the wild": SM

July 5, 2023

**3 Supplementary figures** **6**

### 1 Studying collective behaviour in the wild

Studying collective behaviour in the wild is challenging due to the difficulty of observing animal groups and their interactions. First, it requires us to identify a study system that allows us to observe large groups exhibiting collective movement dynamics. Animal groups exhibiting fission-fusion dynamics are interesting because no obvious hierarchical structure can be easily maintained. Therefore, local rules and emergent properties are more likely to be important in the comprehension of leader-follower rules and predictors of the merge-split outcomes. Moreover, from a logistical point of view, terrestrial animals that live in open habitats are well suited to collective behaviour studies since we can track their movement in two dimensions as opposed to fish schools or bird flocks in three-dimensional spaces spanning large areas. Based on these requirements, we chose our study system, an antelope species - **Blackbuck** (*Antilope cervicapra*).

#### 1.1 Study species: Blackbuck (*Antilope cervicapra*)

Blackbuck are antelopes native to the Indian subcontinent. Blackbuck are social animals, and they form large groups. Group composition varies from all male, and all female to mixed-sex herds (Figure 1-A). These groups exhibit fission-fusion dynamics, and group membership is often dictated by the availability of forage and the nature of the habitat. Males and females differ in body size. Males have long, ringed horns, 35–75 centimetres long. In breeding season, some adult males (Figure 1-A-ii) acquire dark colour, this colour is associated with testosterone levels [Ranjitsinh, 1982]. Blackbuck generally exhibit resource defence polygyny, many males occupy resource territories, and females visit them for mating. Sub-adult males join mobile bachelor herds and sometimes mix in female groups. Blackbuck inhabit grassy plains and slightly forested areas and they forage on short grasses [Jhala & Isvaran, 2016].

##### 1.2 Study site

The study was conducted in Blackbuck National Park, Gujarat. This national park is dedicated to the conservation of blackbuck. It is a semi-arid grassland spread over an area of 34.08 square km. This National Park encloses exclusive grassland habitat, shrublands, saline lands and mud plains. The park embraces over 140 species of birds, 14 species of mammals, 95 species of flowering plants, and many reptiles. The park's fauna includes blackbuck, wolves, hyenas and lesser floricans. The total blackbuck population in and around the park is estimated to be around 4000 (Management Plan for BBNP, 2014-2015).

#### 2 Individual-level response: Traits

We examined the role of trait heterogeneity in governing the response propensity of individuals. Using the methods explained in sections 3.3.1 and 3.3.3, we obtained lists of the initiators, laggers and influential individuals for each video. To maintain rigour, we made separate lists for the top five, top 1/4th, and top 1/3rd individuals appearing as response initiators or leaders. We then made trait tables listing each relevant individual's age/sex categories and spatial positions (before perturbation and during collective motion). Therefore, our conclusions remain consistent irrespective of our threshold to classify response-initiators, isolated and influential individuals.

##### 2.1 Age/sex categories

By utilizing high-resolution videos, we can zoom in and scrutinize individuals closely, enabling us to distinguish their physical characteristics such as body shape, colour, and horns. With these parameters in mind, we categorize individuals into one of four categories:

1. Breeding males (BM) are characterized as dark-coloured individuals with large horns. In blackbucks, the dark coat colour is an indicator of high testosterone levels, and breeding males exhibit this colouration during the breeding season.

2. Females (F) are distinguished by their slender body shape, light brown coat, and lack of horns.

3. The "Other males" category (M) includes non-breeding males and sub-adults. While non-breeding males may possess fully developed horns, they do not display the dark/black coat colour of breeding males, instead having a brown coat colour. Sub-adults also have shorter horns and brown or light brown coat colour.

4. Juveniles and other individuals (Juv) without clearly discernible body shapes or small horns fall into this category, as they are difficult to place in any other category.

Figure 2-A provides an illustrative example of an image zoomed in from a video, featuring individuals from different age and sex groups. Further, we classified the individuals into their respective age and sex categories across all 26 videos and plotted the distribution of each group within these videos (Figure 2-A).

##### 2.2 Spatial dynamics

As previously described, the experiment involved simulating a predation threat by having an experimenter approach a group from a minimum distance of 200 meters, while an observer recorded the group's behavior. Since we anticipated that individuals would react differently based on their proximity to the approaching experimenter, we divided the group into three distinct regions based on their distance from the experimenter *in the direction of approach*: Near, Mid, and Far. To achieve this, we selected three equal-width stripes (as shown in Figure 2-B) in the frame just before the escape phase began. We then tallied the number of individuals present in each of the three regions across all the videos to create a distribution, which is illustrated in Figure 2-B. This normalized frequency distribution serves as the baseline for the number of individuals present in the near, mid, and far regions *before* the perturbation event.

Once the group begins moving away, we visually divide the group into three regions based on their position *in the direction of group movement*: Front, Mid, and Back. To achieve this, we select three equal-width stripes (as shown in Figure 2-C) in the frame towards the mid of the collective motion phase. We then count the number of individuals present in each of the three regions across all the videos and create a normalized frequency distribution, represented as a normalized histogram in Figure 2-C. This normalized frequency distribution serves as the baseline for the number of individuals present in the front, mid, and back regions of the group *during* movement.

#### 2.3 Statistical Methods

##### 2.3.1 Relative role of the attraction and alignment tendencies

Our goal was to find out if the mean $C_{max}$ values and mean lag at $C_{max}$ for any given pair between polarization (m), median NND (d) and median speed (s) were significantly greater than zero or not (SM Figure 3). Since the data did not follow a normal distribution, we carried out non-parametric bootstrapping where sample data was re-sampled with replacement 1000 times, and the mean of these 1000 samples was calculated. We then calculated the 95% confidence interval around the mean. If the interval between values with probability = 0.025 and 0.975 did not contain zero, then the sample statistic was statistically discriminable from zero at an alpha of 5%.

##### 2.3.2 Significance of traits impacting propensity towards individual behaviour

Each trait table is a combination of the following sets: categories (age-sex category, spatial position before perturbation, spatial position during collective motion), individual response (initiators, influential individuals, isolated individuals) and cut-offs (top 5, top 1/4th, top 1/3rd). The statistical results reported below were conducted for the "top 5" threshold for classifying initiators and influential individuals. However, it must be noted that the representation of traits within the response categories remained comparable for all thresholds.

The mean proportion of individuals in each category (for example, the representation of Breeding males in the initiator category) was calculated. Based on this, our null hypothesis for a case where we want to check if the spatial position before perturbation had any impact on initiation would be:

$$\mu_{initiator,Near} = \mu_{initiator,Far} = \mu_{intiator,Mid} \quad (1)$$

We carried out a Kruskal-Wallis test, which is a non-parametric variant of the one-way ANOVA. Since we considered a 95% CI for the tests, if our obtained p-value was $> 0.05$, we had to reject our null hypothesis. We carried out a pairwise Wilcoxon test with a Benjamini-Hochberg p-value correction in order to find the pairs whose mean proportion of individuals differed significantly between the categories in each dataset. Pairs with a significant difference in their means had a $p - value < 0.05$.

Details of the statistical tests and results:

1. **Friedman Test to check for differences in network connectedness between response phases (Figure 3-A in main text)**: null hypothesis = no difference in mean of Proportion of unconnected nodes between any phases.
   Friedman is the non-parametric test for repeated measures ANOVA for three or more paired groups).
   Effect size: 0.268, small
   p-value=0.0051
   CI = 95% or alpha = 0.05
   p-value < 0.05. Null hypothesis rejected.

2. **Wilcoxon signed rank test**: Post-hoc test to find the pairs of response phases with the significant differences between network connectedness. Since, we have ${}^4C_2$ pairs, we need to adjust p-values to account for these ${}^nC_2$ pairwise test. We did a Bonferroni adjustment. p-vales between the pairs are given below.

Ini-esc:0.0015
Ini-cmp: 0.0609
Ini-Relaxation: 0.6152
Esc-cmp: 0.7595
cmp-relaxation: 1.00
esc-relaxation: 0.3140

Only ini-esc p-value $< 0.05$. Therefore, it is the only significantly different pair. Meaning, there is a significant difference in the network connectedness only between the initial unperturbed phase to the escape phase.

3. **Influence of Age/sex categories:**
   Kruskal-Wallis Test, p-values given below: CI=95%
   Influential individuals =0.6362
   Isolated = 0.9774
   Initiators = 0.3052
   all p-vales $> 0.05$, therefore, no significant differences. No need for post-hoc test.

4. **Influence of Positions (before perturbation):**
   Kruskal-Wallis Test, p-values given below: Influential individuals: p-value = 0.4679
   Isolated: p-value = $0.02185 < 0.05$
   Effect size = 0.0724: medium

   Wilcoxon rank sum test: post-hoc test.
   p-value adjustment = BH-method.
   p-values:
   Near-mid: 0.982
   mid-far: 0.031
   neaf-far: 0.031
   Therefore, isolated individuals are significantly more likely to be found in the far region.

   Initiators:
   p-value = $0.004178 < 0.05$ (Kruskal-Wallis Test)
   Effect size = 0.0558: small
   Wilcoxon rank sum test: post-hoc test.
   p-value adjustment = BH-method.
   Near-mid: 0.7250
   mid-far: 0.077
   near-far: 0.0271
   Therefore, initiators are significantly less likely to be found in the far region.

5. **Positions (during collective movement):**
   Leaders: p-value for Kruskal-Wallis = 0.0036 ¡ 0.05
   Effect size = 0.0629: medium

   Wilcoxon rank sum test: post-hoc test. p-value adjustment = BH-method. p-values:
   Front-mid: 0.029
   mid-back: 0.321
   front-back: 0.004
   Therefore, leaders are significantly less likely to be found in the front of the group.

Isolated: p-value for Kruskal-Wallis = 0.0016 ¡ 0.05
Effect size = 0.0626: medium
Wilcoxon rank sum test: post-hoc test. p-value adjustment = BH-method. p-values:
Front-mid: 0.0033
mid-back: 0.6970
front-back:0.0033
Therefore, isolated individuals are significantly less likely to be found in the front of the group.

**Summary of statistical analysis results:**

1. Age-sex categories: not statistically significant/discriminable
2. Distance from threat (before perturbation):
   Leader: not statistically significant/discriminable.
   Solitary: Near and far, mid and far. i.e. far is significantly (statistically) different (highest proportion).
   Initiators: Near and far, mid and far. i.e. far is significantly (statistically) different (lowest proportion).
3. Position in the group (Collective motion):
   Leaders: front and mid, front and back, i.e. front is significantly (statistically) different (lowest proportion).
   Solitary: front and mid, front and back, i.e. front is significantly (statistically) different (lowest proportion).
   Initiators: not statistically significant/discriminable.

Based on the results Age and Sex doesn't seem to be playing a role in determining which individual identifies the threat first or has the most influence on the group movement during the collective motion phase.

However, the individuals who are farthest away from the threat during the approach are least likely to be the escape initiators and interestingly these individuals later on have the least correlated movement with other group members. We postulate that such group members might have a possibility of splitting away from the group.

During the collective motion phase, individuals at the front are least likely to influence the speed of other individuals. This might be due to a push-like drive from the individuals at the back rather than trailing individuals trying to catch up with the front individuals. As expected, movement initiators do not occupy any typical position in the group during the collective motion.

[Ranjitsinh, 1982] Ranjitsinh, MK. 1982. Territorial behaviour of the Indian Blackbuck (*Antilope Cervicapra*, LINNACUS, 1758) in the Velavadar National Park, Gujarat.

[Rathore, 2021] Rathore, Akanksha. 2021. *Collective escape dynamics of fission-fusion groups in the wild.* Ph.D. thesis.

### 3 Supplementary figures

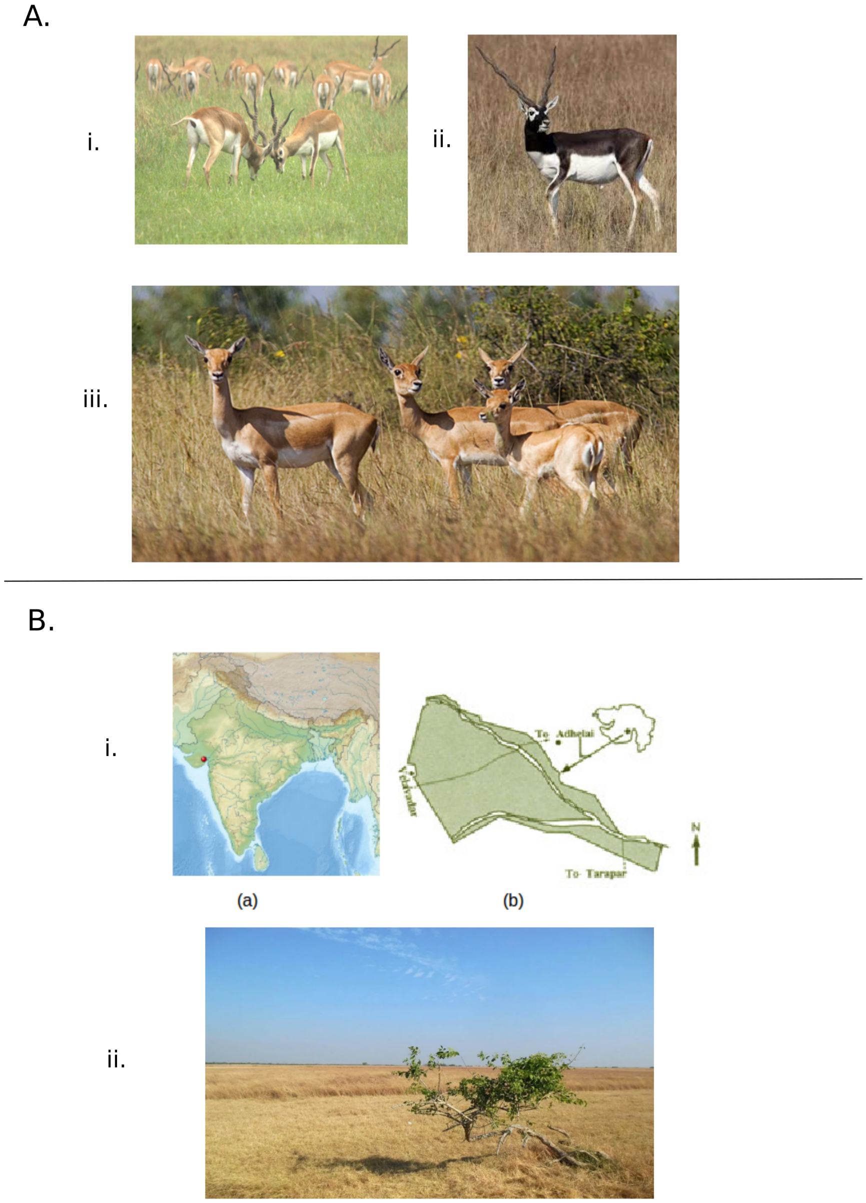

Figure 1: A. Blackbuck males and females are dimorphic, varying in body size, colour and presence of horns. A.i. Two sub-adult males engaged in a tactile fight. A.ii. A breeding male - males acquire dark-coloured coats in the breeding season. A.iii. A herd of adult females and juveniles. B.i. Blackbuck National Park, Velavadar (Gujarat). (a) Location on Indian map, (b) Geographic map of the national park ( Pic credits: Gujarat forest department website). B.ii. This park is majorly a semi-arid grassland. Figure from [Rathore, 2021] PhD thesis.

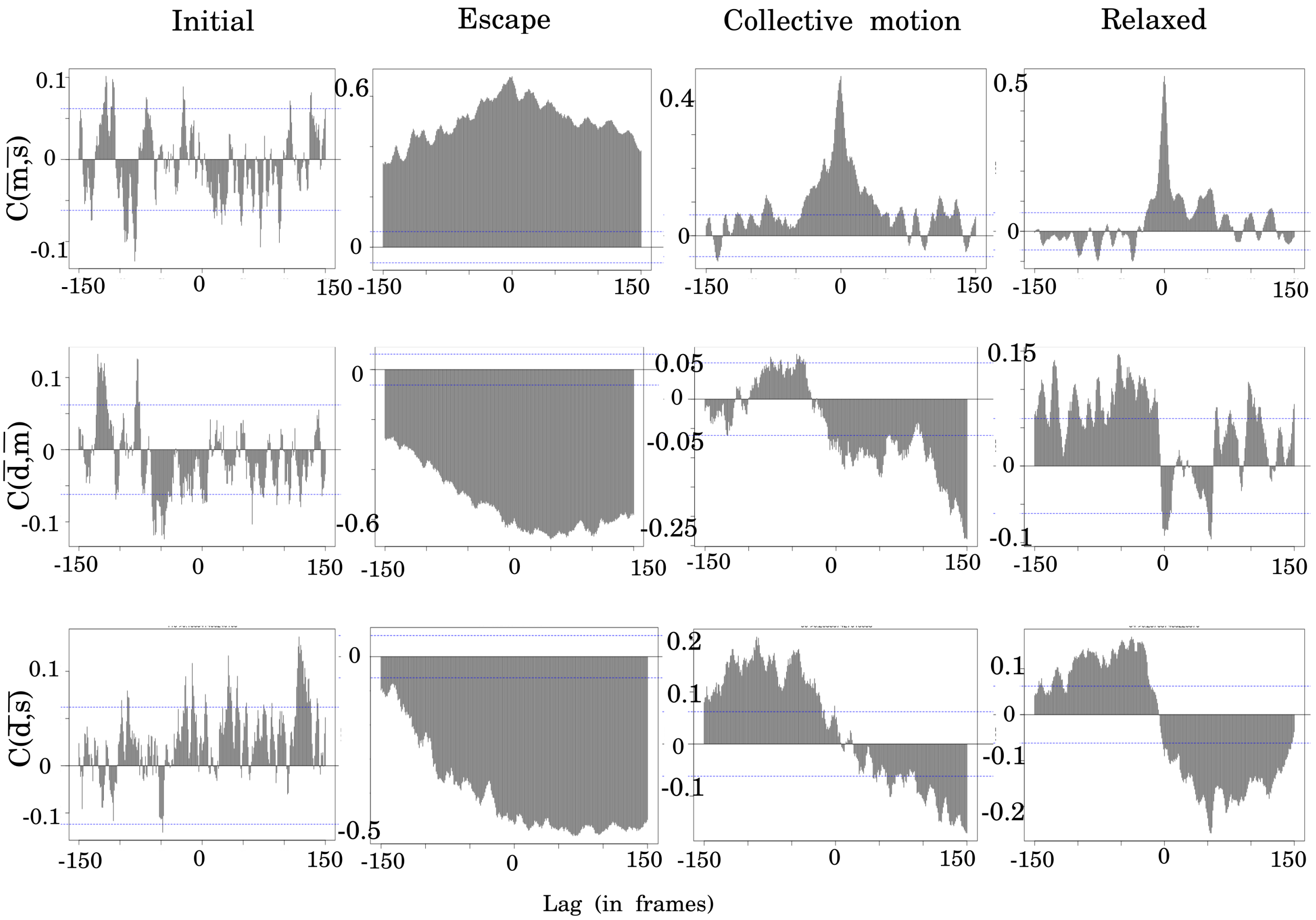

Figure 2: Cross-correlation output for the four phases of an example video for all three pairs of the movement metrics – i. $C(\overline{m}, \overline{s})$: Polarization and median speed of the individuals; ii. $C(\overline{d}, \overline{m})$: median NND and polarization; iii. $C(\overline{d}, \overline{s})$: median NND and median speed of the individuals. We observe a change in the sign and as well as value for the maximum correlation strength and lag for the escape and collective motion phases. The values for the initial and relaxation phases are similar.

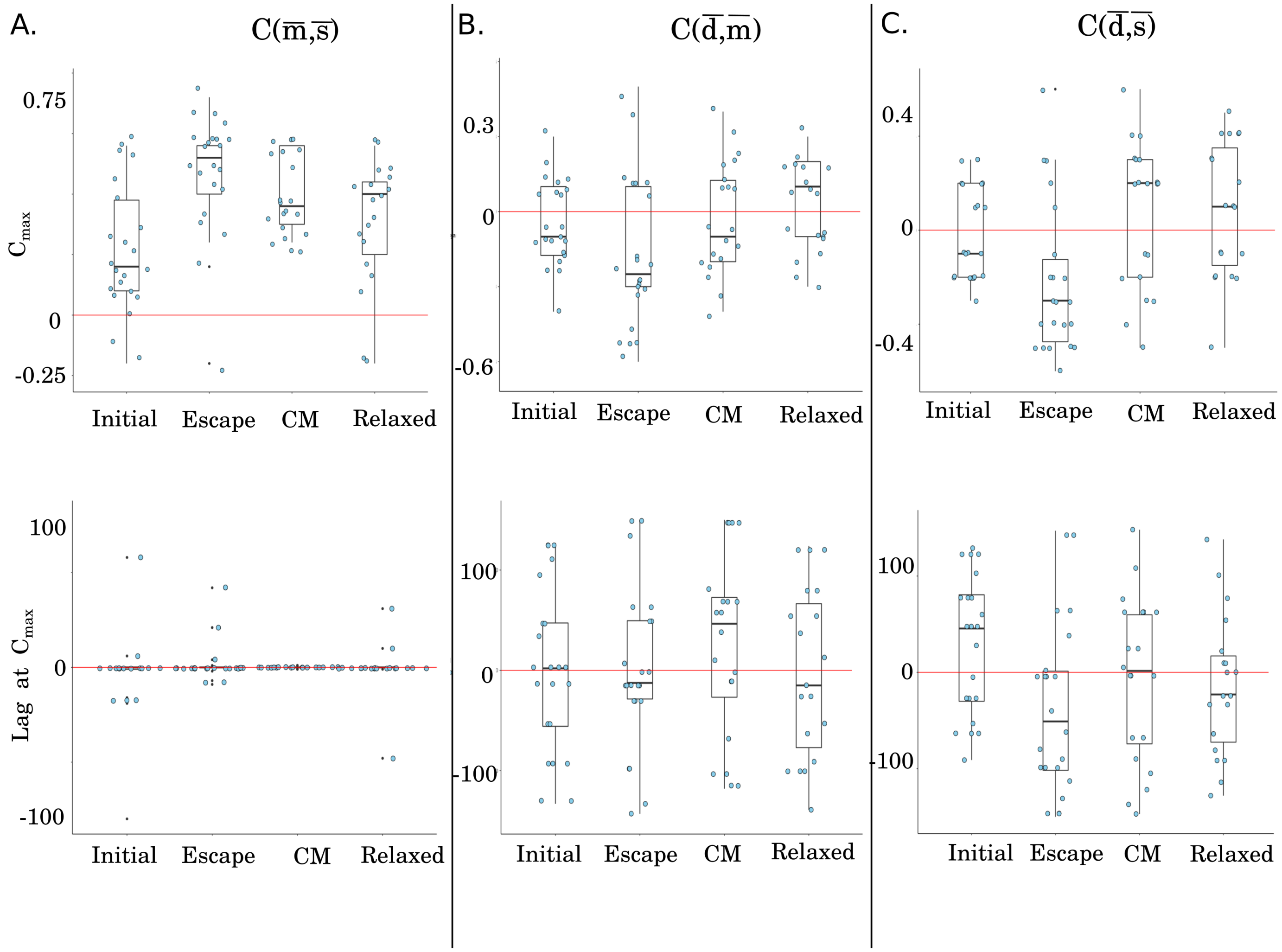

Figure 3: Cross-correlation output for the four phases across all videos for all three pairs of the movement metrics – A. $C(\overline{m}, \overline{s})$: Polarization and median speed of the individuals; B. $C(\overline{d}, \overline{m})$: median NND and polarization; C. $C(\overline{d}, \overline{s})$: median NND and median speed of the individuals. We observe a change in the sign and as well as value for the maximum correlation strength and lag for the escape and collective motion phases. The values for the initial and relaxation phases are similar.

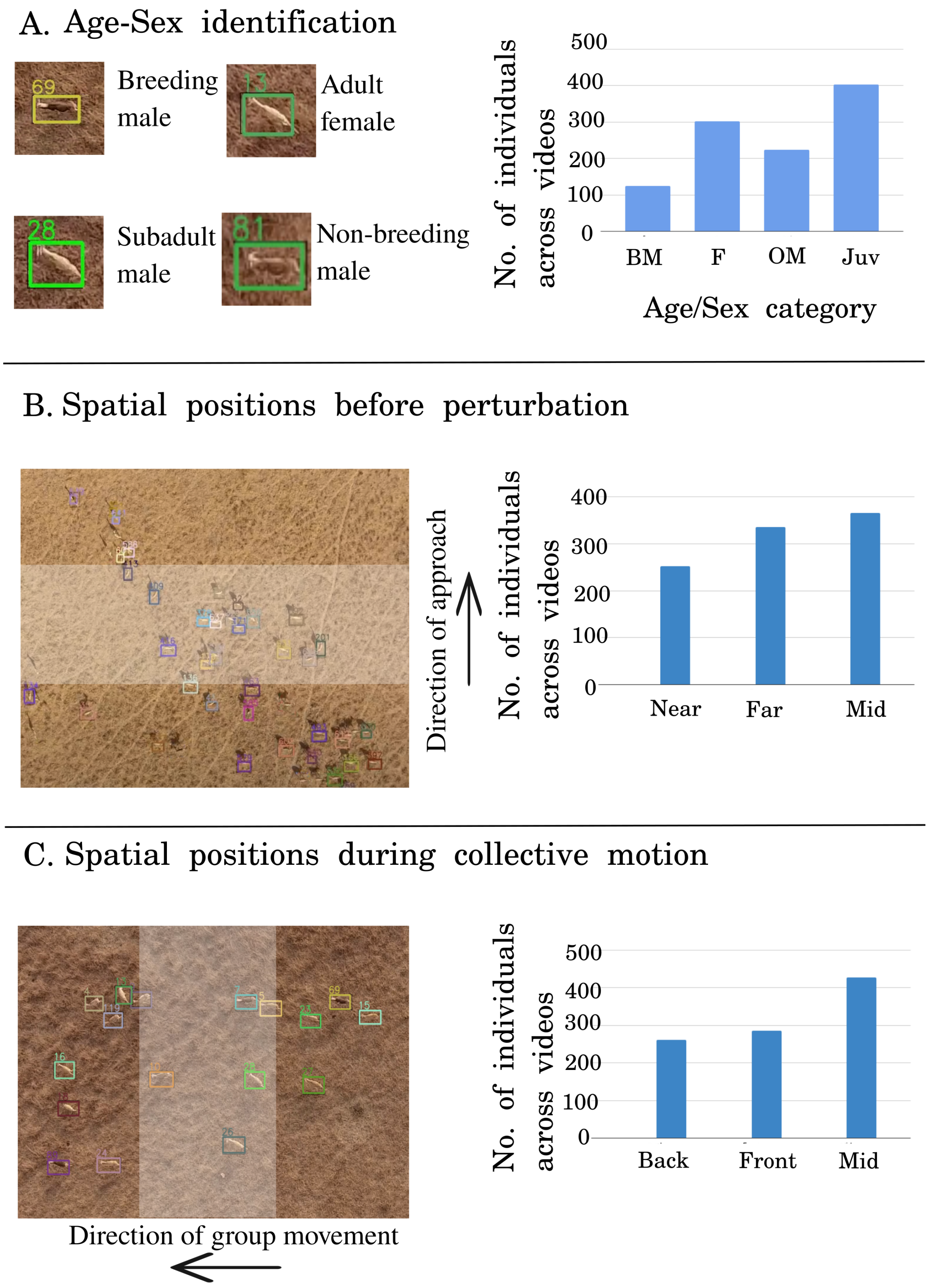

Figure 4: A. Zoomed-in image from a video. Individuals of different age/sex classes are visible in this frame. The right panel shows the distribution of various categories across all the videos. B. Spatial distribution of individuals at different distances from threat, the right panel shows the frequency histogram in the three categories - near, mid, and far. C. Spatial distribution of individuals during the group movement, the right panel shows the frequency histogram in the three categories - front, mid, and back.

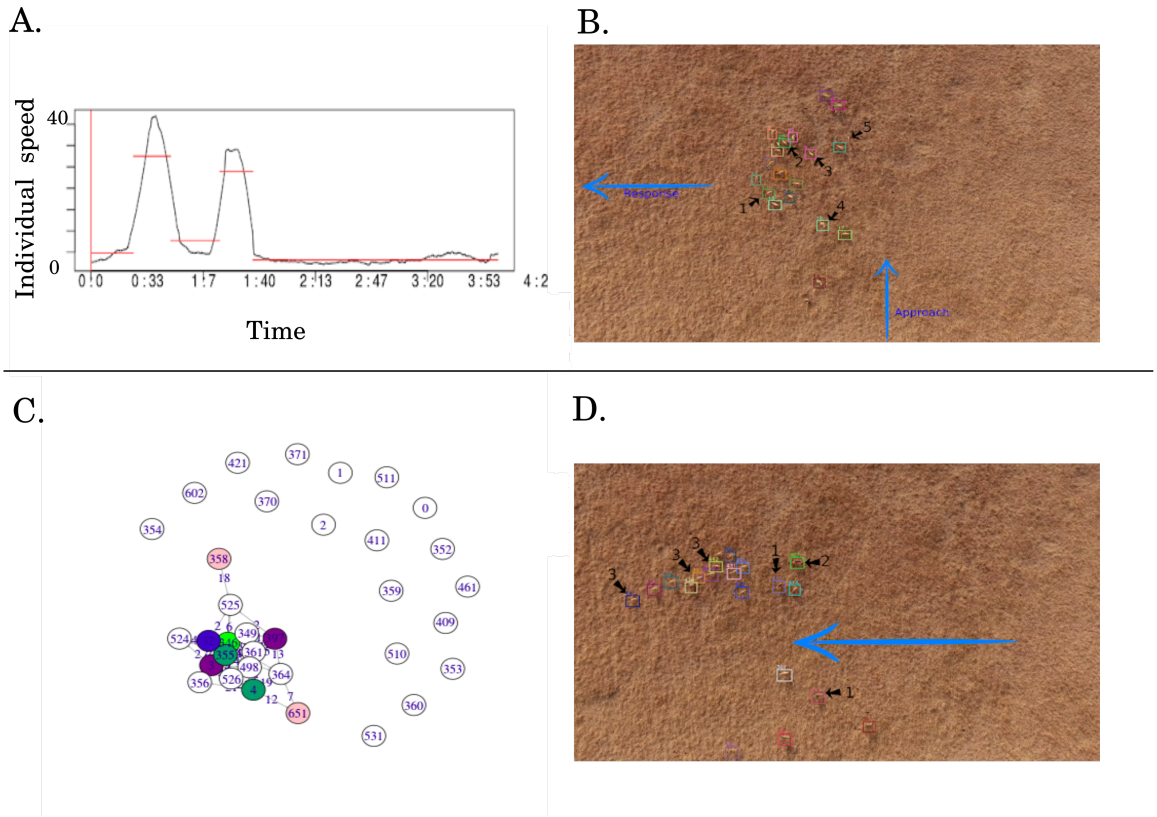

Figure 5: A. Identifying the response initiators: Smoothened time series of speed for one individual where red lines show different stages as determined by the change point method. (B) Initiators were identified using change point analysis of speed time series, plotted on one frame of the video. C. Identifying the influential individuals/leaders: The nodes of the network colour coded for the number of connections with a minimum time lag. D. Top five ranking individuals mapped on a frame of the example video.

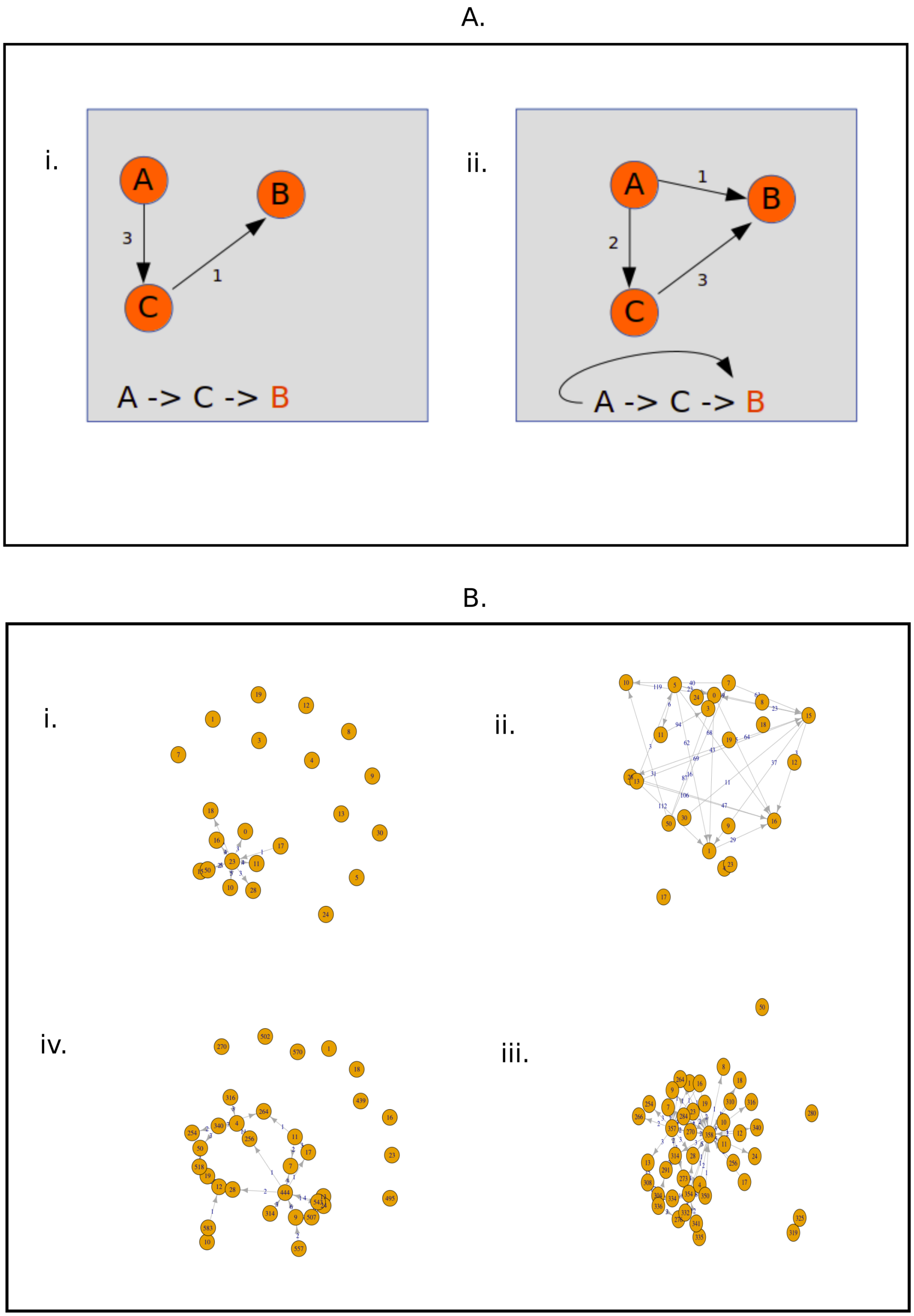

Figure 6: Leader-follower network built using cross-correlation functions between individual speed time series. (A) Two example scenarios of lead-lag relationships between individuals in a group of three members. (B) Lead-lag network of individual speeds for one sample video, i. Initial state - before the approach starts, ii. Escape phase - when individuals start responding to the approach, iii. Coordinated bout - network when all individuals are moving together in a coordinated way, iv. Post response - network when individuals have stopped moving and are back to their foraging/resting activities.

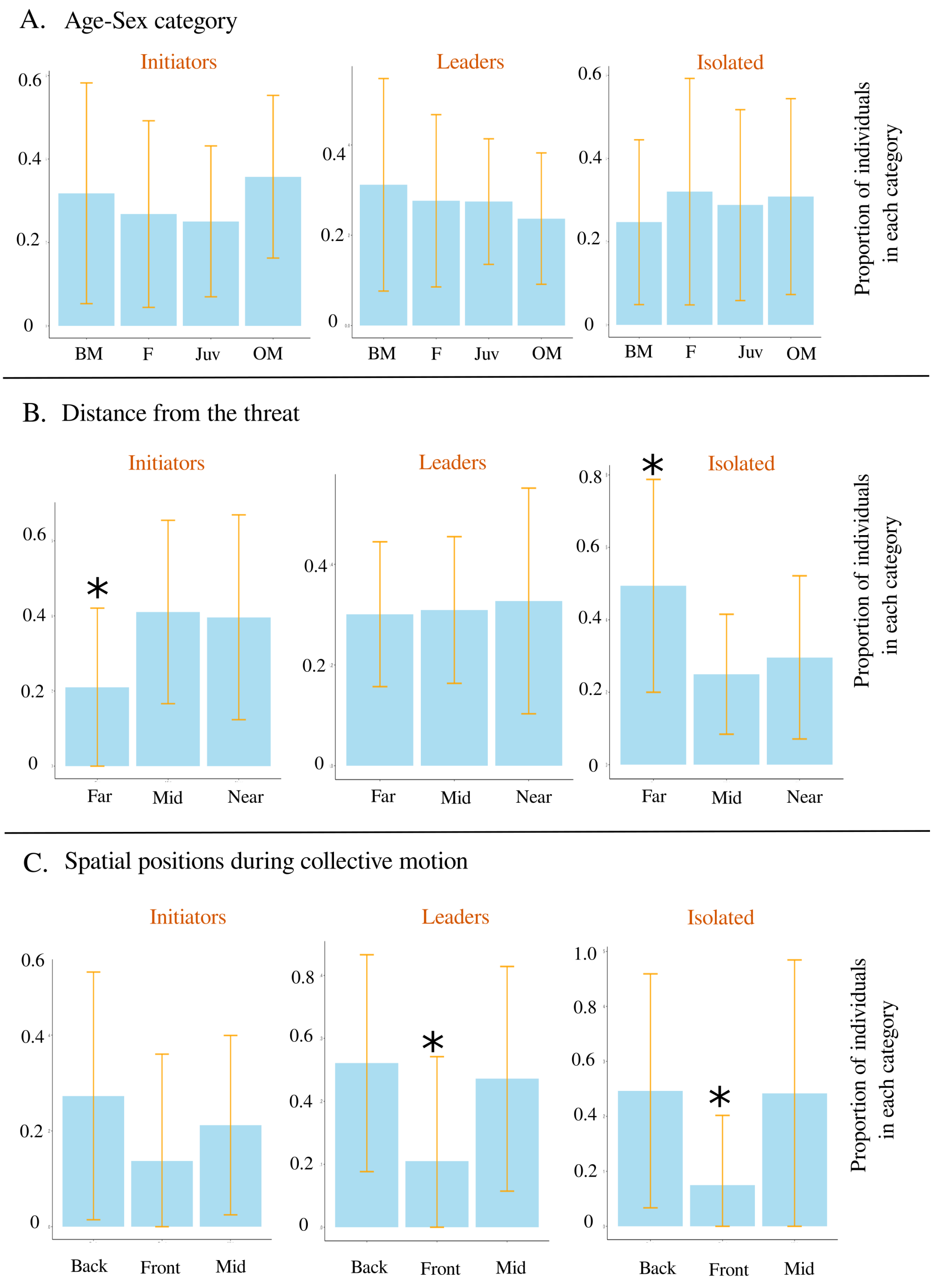

Figure 7: A. The distribution of individuals initiating the escape response, exerting the greatest influence on group movement, and whose speed is uncorrelated with group movement based on their age/sex category. B. The distribution of individuals initiating the escape response, exerting the greatest influence on group movement, and whose speed is uncorrelated with group movement based on their distance from the approach. C. The distribution of individuals initiating the escape response, exerting the greatest influence on group movement, and whose speed is uncorrelated with group movement based on their position in the group w.r.t the direction of group movement. The * mark represents the category for which the differences in pair-wise comparisons are statistically significant after non-parametric bootstrap sampling.
